# Maternal malaria and early-life infections shape humoral immunity to *P. falciparum* in infants

**DOI:** 10.64898/2026.02.13.705422

**Authors:** Felistas Nankya Namirimu, Florian A. Bach, Kathleen Dantzler Press, Pamela Odorizzi, Kenneth Musinguzi, Kate Naluwu, Isaac Ssewanyana, Abel Kakuru, Moses Kamya, Grant Dorsey, Margaret E. Feeney, Prasanna Jagannathan

**Affiliations:** Infectious Diseases Research Collaboration, Kampala, Uganda; Department of Medicine, Stanford University, Stanford, CA, United States; University of California San Francisco, San Francisco, CA, United States; Gilead Sciences, United States

## Abstract

Placental malaria may negatively impact acquisition of immunity to malaria in infants. To study the effect of maternal antibodies on early-life infections, a birth cohort of infants were enrolled in a prospective observational study in Busia, Uganda. We measured *P. falciparum*-specific antibody responses to 19 *P. falciparum* antigens and tetanus toxoid in 183 infants using a Luminex multiplex bead array. We also assessed the frequency and phenotype of peripheral T follicular helper cells and B cells using flow cytometry. Placental malaria and lower gestational age adversely impacted antibody transfer overall. Maternal antibodies present in cord blood were not associated with protection in the first year of life, and neither was maternal malaria status. Infants generated antibodies against *P. falciparum* antigens in response to infection, but these were also not associated with protection, only exposure. Interestingly, infections in the first six months of life correlated with decreased levels of antibodies to *P. falciparum* antigen at one year of age. Early-life infections were not clearly associated with circulating follicular T cell or B cell phenotypic composition. These data suggest the infant antibody response has little, if any, impact on preventing clinical malaria or patent parasitemia in the first year of life.

## Introduction

Malaria remains one of the most severe public health problems worldwide with an estimated 263 million cases and 597,000 deaths^1^. The World Health Organization recommends implementing prevention strategies based on vector control, prompt diagnosis and treatment of malaria, chemoprevention, and vaccines^2^. Despite these control interventions, a large burden of malaria persists in endemic settings. Moreover, acquisition of resistance phenotypes by *Plasmodium spp.* parasites^3^ and *Anopheles spp.* vectors^4^ put the toolbox against malaria at risk of losing efficacy. Careful study of how protective immunity to malaria is acquired is therefore timely if we are to create new and better interventions.

Immunity to malaria critically depends on both cell-mediated and humoral (antibody) responses. Functions of the latter include complement fixation, invasion inhibition, antibody-dependent cellular phagocytosis and antibody-dependent cellular cytotoxicity. Though individual antibody specificities or effector functions provide only incremental utility, breadth and depth of the antibody repertoire is associated with protection^5,6^. Antibody responses are shaped by T follicular helper (Tfh) cells that provide crucial help for B cells in the germinal center, enabling them center to undergo activation, affinity maturation, class-switching, and differentiation into memory B cells and antibody-secreting plasma cells. The differentiation of B cells has been reported to be negatively impacted by malaria, chiefly due to the expansion of atypical B cells^7^, which may exhibit impaired B cell receptor signaling and effector function^8^. In contrast, atypical B cells are also associated with increased breadth of humoral responses^9^ and a source of neutralizing antibodies specific to *P. vivax*^10^. They exhibit somatic mutation and class switching and expand robustly following vaccination with whole *P. falciparum* sporozoites as well as the inactivated influenza vaccine^11^. This uncertainty about whether atypical B cells impact malaria protection positively or negatively (or either, depending on the context) and their potential for eliciting effective immunity warrants further study of their occurrence in at-risk populations.

In high-transmission settings, first malaria infections typically occur in the first year of life, when pre-existing, maternally-transferred antibodies coexist with antibodies made by the infant in response to infection. While maternal antibodies may be protective, they may also hamper the development of immunity in the infant, for example due to epitope masking in the germinal center^12^. Adult and infant immunoglobulins cannot be readily distinguished molecularly, meaning it is unclear how long malaria-specific maternal antibodies may persist, and how they may contribute to emerging immunity in the infant.

In this study we present a unique birth cohort of children born in a high-transmission setting in Eastern Uganda. Due to efficacious chemoprevention and vector control, malaria incidence in the first year of life was low (∼ 0.2 episodes per person year), meaning that the kinetics of malaria-specific maternal antibodies could be determined, with limited confounding from infant immune responses.

## Results

### Characteristics of study participants

A total of 186 children were included in the study. Plasma was collected from cord blood at birth and venous blood at 6 and 12 months. There was no association between maternal chemoprevention treatment group and antibody levels in cord blood (**Fig. S1**), so results were pooled for all subsequent analyses. A custom Magpix bead assay was used to measure antibodies to 20 targets: 19 *P. falciparum* antigens as well as tetanus toxoid. These antigens were chosen to cover a range of parasite subcellular localization and function, good markers of exposure, and candidate markers for protection. Fresh peripheral blood mononuclear cells from partially overlapping groups of 99 and 93 children were collected for T and B cell phenotyping by flow cytometry, respectively. All three assays were available for 54 children (**Fig. S2**). Demographic characteristics are summarized in **Table 2**.

**Table 1:** Characteristics of study participants.

|  | Clinical Cohort (186 children born to 183 mothers) |
| --- | --- |
| **Maternal characteristics** |  |
| Age (years) - median (IQR) | 21.9 (19.0 — 24.77) |
| Gravidity – n (%) |  |
| 1 | 54 (29.5%) |
| 2 | 62 (33.9%) |
| ≥3 | 67 (36.6%) |
| IPT arm – n (%) |  |
| 3 Dose SP | 97 (53.4%) |
| 3 Dose DP | 42 (23.8%) |
| Monthly DP | 44 (23.8%) |
| **Maternal exposure during pregnancy** |  |
| Malaria Exposure Categories – n (%) |  |
| No malaria in pregnancy | 93 (50.8%) |
| Non-placental malaria | 17 (9.3%) |
| Placental malaria | 69 (37.7%) |
| Unknown | 4 (2.2%) |
| **Infant characteristics at birth** |  |
| Gestational age (weeks) – median (IQR) | 39.6 (38.9-40.7) |
| Low birth weight (<2500g) – n (%) | 20 (10.8%) |
| Preterm birth (<<37 weeks) – n (%) | 12 (6.4%) |
| Gender (Female) – n (%) | 93 (50%) |
| IPTc arm – n (%) |  |
| DP every 12 weeks | 185 (99.5%) |
| DP every 4 weeks | 1 (0.5%) |

Abbreviations: IQR: Interquartile Range, IPTp: Intermittent Preventive Treatment during Pregnancy, IPTc: Intermittent Preventive Treatment during Childhood, DP: Dihydroartemisinin Piperazine, SP: Sulfadoxine Pyrimethamine

**Table 2:** Summary of the P. falciparum Blood stage antigens.

| Gene ID | Antigen Name | Abbreviation |
| --- | --- | --- |
| PF3D7_1133400 | Apical Membrane Antigen 1 | AMA1 |
| PF3D7_0304600 | Circumsporozoite Protein | CSP |
| PF3D7_1301600 | Erythrocyte Binding Antigen-140 (R III-V) | EBA140 |
| PF3D7_0731500 | Erythrocyte Binding Antigen-175 (R III-V) | EBA175 |
| PF3D7_0102500 | Erythrocyte Binding Antigen-181 (R III-V) | EBA181 |
| PF3D7_0423700 | Early Transcribed Membrane Protein 4 Antigen 1 | ETRAMP4 |
| PF3D7_0532100 | early transcribed membrane protein 5 Antigen 1 | ETRAMP5 |
| PF3D7_1035300 | Glutamate Rich Protein (R2) | GLURP |
| PF3D7_1036000 | Merozoite Surface Protein 11 | H103 |
| PF3D7_0501100.1 | Heat Shock Protein 40 (Antigen 1) | HSP40 |
| PF3D7_1002000 | Plasmodium exported protein (hyp2) | HYP2 |
| PF3D7_0930300 | Merozoite Surface Protein (19kDa fragment) | MSP1 |
| PF3D7_0206800 | Merozoite surface protein 2 (CH150/9) | MSP2-CH150 |
| PF3D7_0206800 | Merozoite surface protein 2 (DD2 allele) | MSP2-DD2 |
| PF3D7_1335400 | Reticulocyte Binding Protein Homologue 2 | RH2 |
| PF3D7_0424200 | Reticulocyte Binding Protein Homologue 4 | RH4 |
| PF3D7_0424100 | Reticulocyte Binding Protein Homologue 5 | RH5 |
| PF3D7_0501300 | Skeleton-Binding Protein 1 | SBP1 |
| PF3D7_1021800 | Schizont Egress Antigen 1 | SEA |
| n/a | Tetanus Toxoid (Non-adsorbed) | TT |

### Placental malaria and low gestational age negatively impact transfer of antibodies

To investigate whether *in utero* malaria exposure affected antibody transfer, we compared cord blood antibody levels of children stratified by whether their mother had evidence of placental malaria, non-placental malaria, or no malaria in pregnancy at all. 5 of the 20 antibodies were differentially abundant in children born to mothers with placental malaria, compared to no malaria: those binding AMA1, SEA, GEXP, HYP2, and tetanus toxoid (**Fig. 1A**).

**Figure 1:**
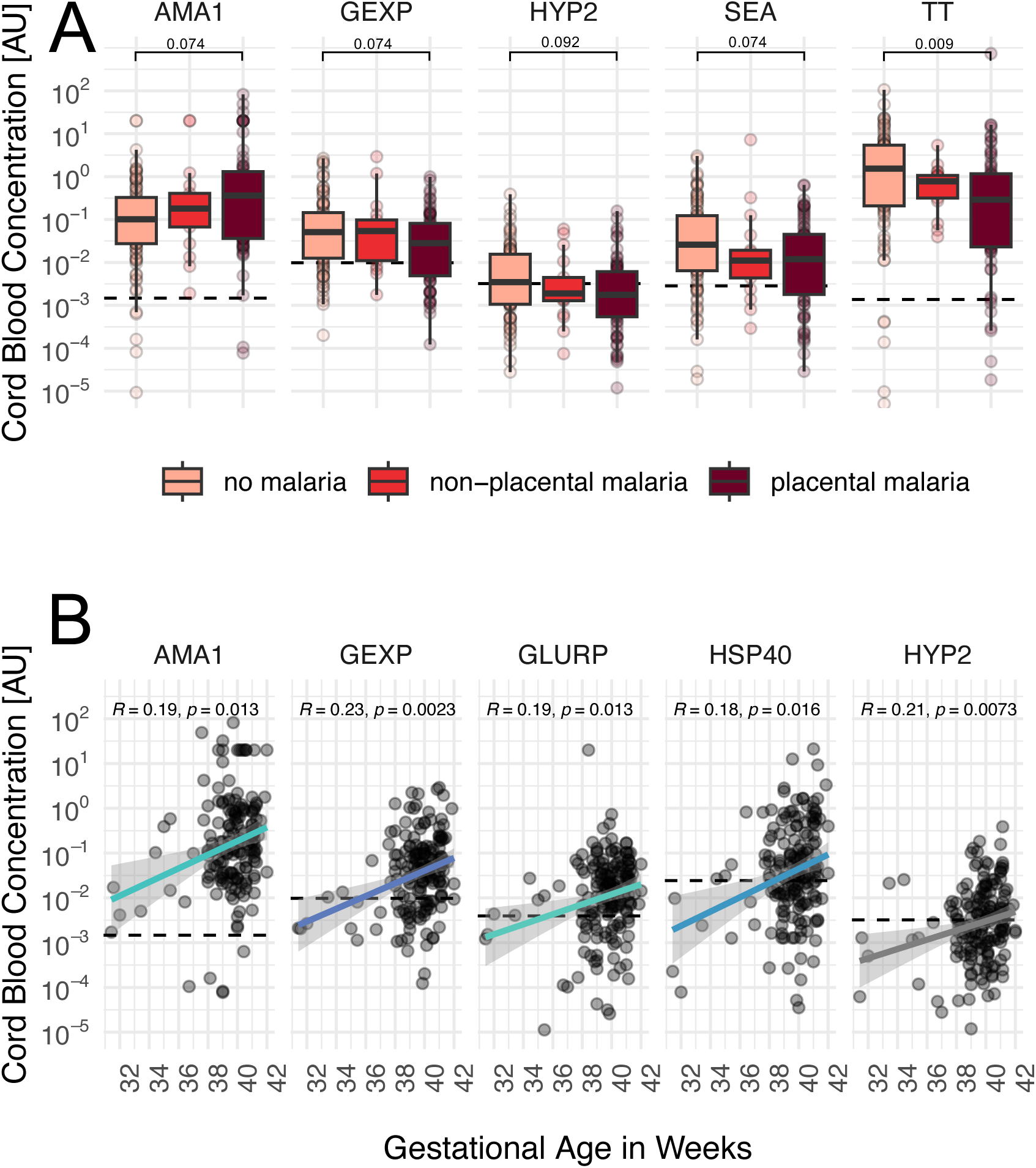
Maternal malaria and pre-term birth negatively impact antibody transfer. **(A)** Antibody concentrations in cord blood are stratified by maternal malaria outcomes during pregnancy. P values were obtained from Wilcoxon signed-rank test, corrected for multiple testing using Benjamini Hochberg. All antigens with adjusted p values < 0.1 are shown. **(B)** Spearman correlations with unadjusted p values between gestational age and antibody concentrations in cord blood. For presentation purposes, not all significant associations are shown. The complete list of associations is shown in Fig. S4.

Antibodies to to *P. falciparum* antigens were generally less abundant in children born to mothers who experienced placental malaria (**Fig. S3**), consistent with some prior reports^13,14^. Responses to tetanus toxoid were also lower in the cord blood of children born to mothers with placental malaria, suggesting that placental malaria negatively impacts IgG transfer generally, rather than in a pathogen or antigen-specific manner. In contrast, antibodies to AMA1 were increased in the cord blood of children born to mothers with placental malaria. Antibodies recognizing AMA1 are among the most abundant Pf-specific antibodies^15,16^ and antigen-specific boosting of maternal antibodies in responses to infection may thus override a defect in overall transfer.

Transfer of IgG to the fetus is highest late in gestation, and it is known that pre-term birth leads to reduced antibody transfer^17^. Also in our cohort, gestational age was associated with higher antibody responses to most antigens at birth, with 9/20 being significantly positively correlated with children born on or after 37 weeks (**Fig. 1B** and **S4**).

### Maternally derived antibodies wane in the first months of life

After quantifying maternal antibody levels in cord blood, we sought to measure the dynamics of these antibodies in the first year of life. Usually this presents a challenge in endemic populations, because antibodies produced by the infant cannot readily be distinguished from those that are maternally derived. However, due to efficacious chemoprevention and indoor residual spraying, malaria incidence in this cohort was low in the first year of life (∼0.2 episodes per person year)^18^, meaning that antibodies present in children who never got infected are most likely of maternal origin. We thus compared antibody concentrations at 6 and 12 months to those measured at birth in a subset of 161 children where neither clinical malaria nor asymptomatic parasite carriage were detected in the first year of life. In the absence of overt infection, antibodies to malaria quickly waned below the seropositivity threshold, suggesting that recent infection is necessary to sustain antibody levels **(Fig. 2)**.

**Figure 2:**
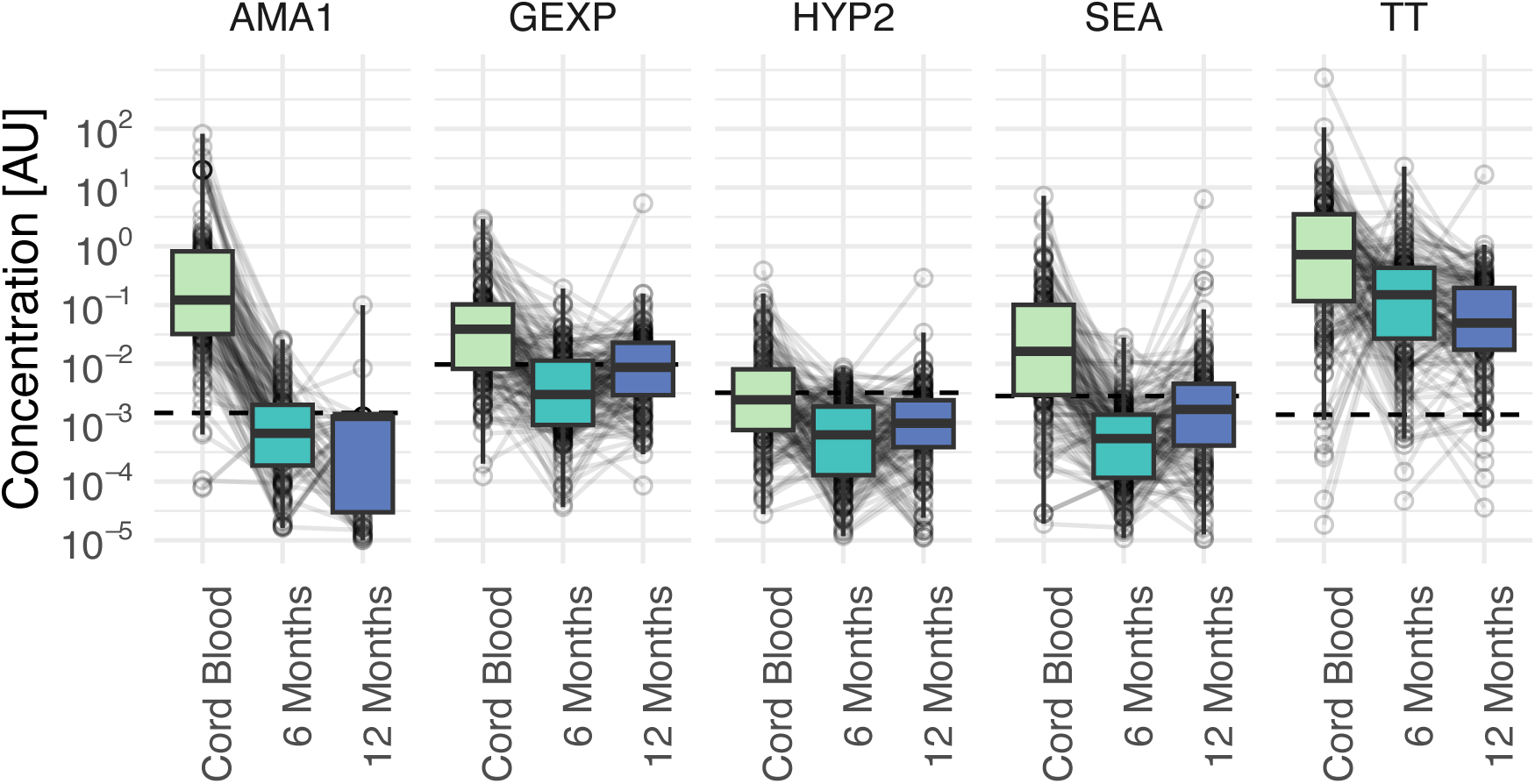
Maternally derived antibodies to *P. falciparum* wane quickly in the first 6 months of life in the absence of infection. Antibody concentrations are shown at different timepoints in the first year of life. For illustration purposes only five analytes are shown, the full list can be seen in Fig. S5.

We saw a small increase in reactivity to most antigens at 12 months compared to 6 months, even in the absence of infection, but the median concentration remained below the seropositivity threshold for the majority of antibodies **(Fig. S5)**. The rapid decay in antibody concentration within 6 months suggests that protection mediated by maternal antibodies, if any, may be most apparent in the first few months of life.

### Cord blood antibodies are not associated with infection incidence in early life

To study the association between infection incidence and cord blood antibodies, we used Poisson regression to model the number of prevalent parasitemic episodes (i.e. the number of months where parasitemia was observed), as well as the number of malaria episodes in the first six months of life. None of the antibodies tested were significantly associated with the incidence in the first six months of life (**Fig. 3**).

**Figure 3:**
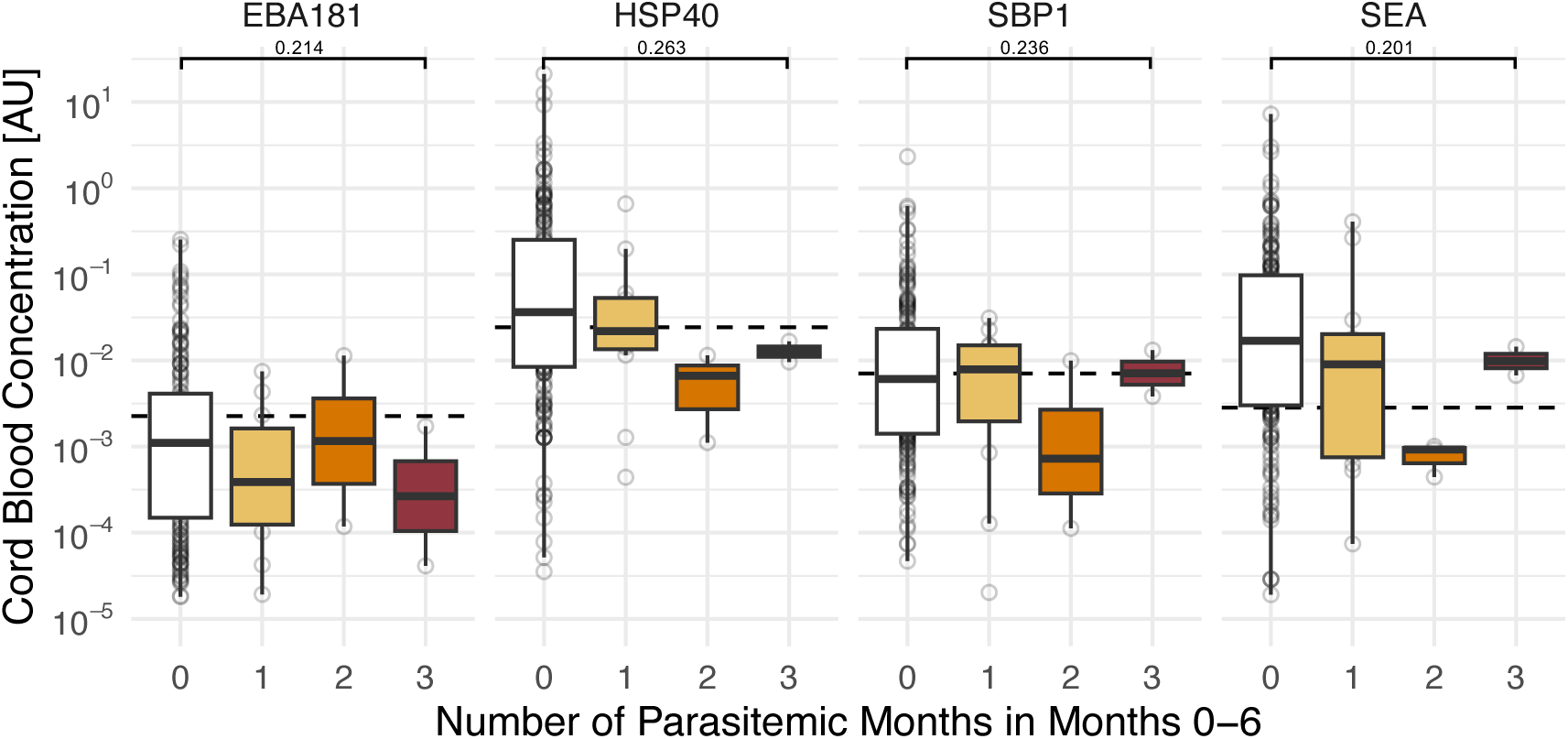
Cord blood antibody levels to *P. falciparum* antigens are not predictive of incidence during infancy. Data are stratified by the number of prevalent infections, i.e. parasitemic months experienced in the first six months of life. P values shown are BH-adjusted p values of Poisson regression coefficients.

### Infant-derived antibodies are markers of recent exposure but are not associated with protection

We next assessed the role of infant antibodies at later timepoints, after maternal antibodies have waned. To this end, we investigated associations of antibody levels at 6 and 12 months with infection prevalence before those timepoints, to identify markers of exposure, and after, to identify markers of protection. We found antibodies to 7 *P. falciparum* proteins (ETRAMP4, ETRAMP5, GLURP, HSP40, HYP2, RH5, SBP1) were higher at 6 months in children who had more parasitemic months in the first 6 months of life (**Fig. 4A**). Likewise, antibodies to 3 *P. falciparum* proteins (ETRAMP5, HYP2 and MSP1) were significantly increased at 12 months in children who got infected between months 6 and 12 **(Fig. 4B).** This suggests that infants can produce IgG recognizing a broad range of *P. falciparum* antigens in response to infection and identifies these antibodies as markers of recent exposure in young infant populations.

**Figure 4:**
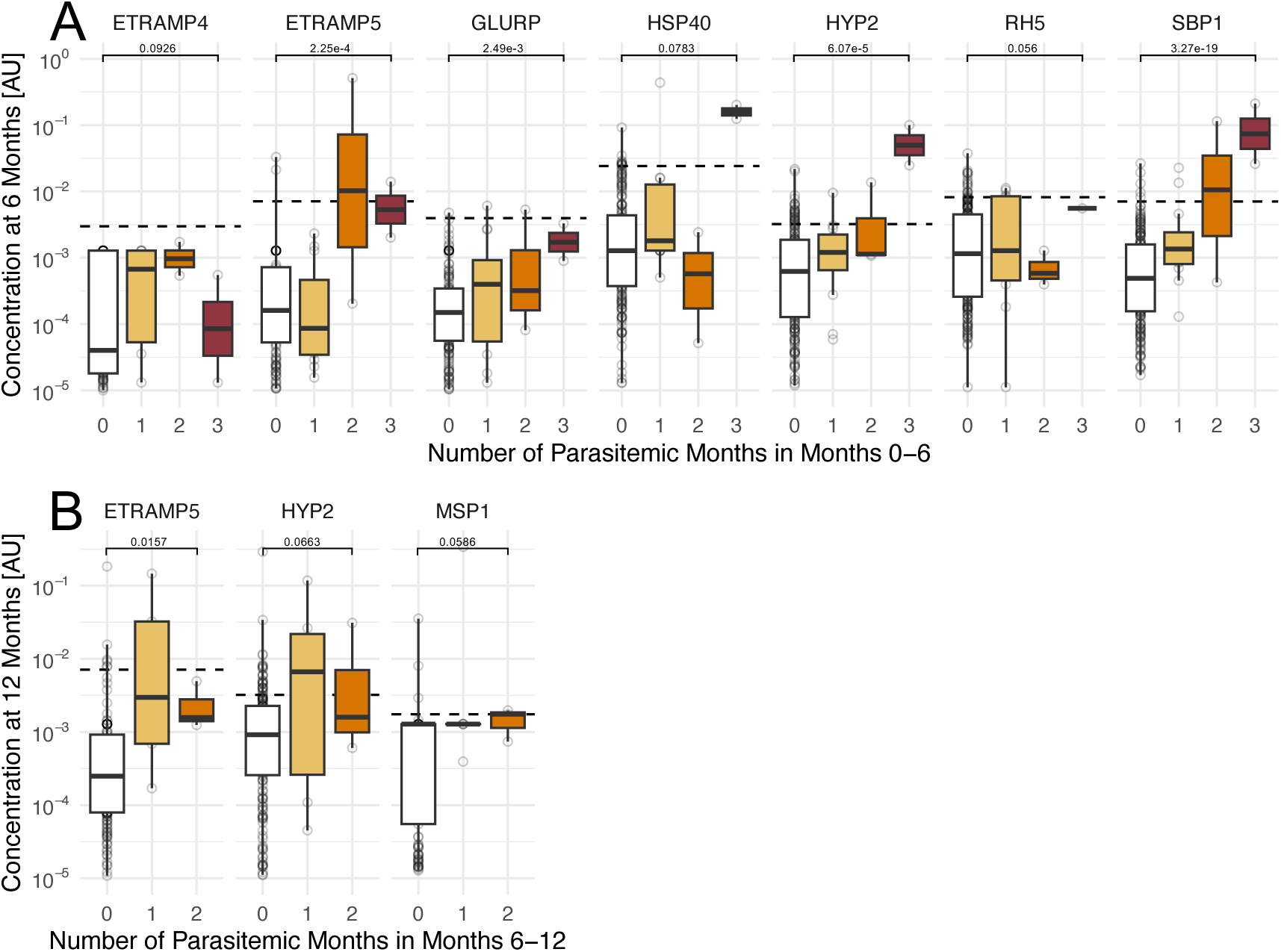
Humoral markers of *P. falciparum* exposure in infants. Antibody concentrations at 6 months (A) and 12 months (B) were significantly associated with increased infections in the 6 months prior to measurement. P values shown are BH-adjusted p values of Poisson regression coefficients.

Strikingly, no antibodies we measured were associated with protection at 6 months or 12 months. To test whether the breadth of the antibody repertoire, rather than individual antibodies themselves, were associated with protection, we calculated a breadth score akin to Nkumama et al.^6^. For each antigen, we sorted individuals into quartiles, based on antibody concentration. The quartile positions (1 to 4) were then summed to calculate a breadth score for each individual and each timepoint. Consistent with the antigen-level antibody responses, the aggregate breadth score was not significantly associated with protection at any of the timepoints measured (**Fig. 5A**). Instead, high breadth was associated with a larger number of prevalent parasitemic episodes in the preceding six month period, though this was only significant at 6 months of age. In addition, we investigated whether placental malaria, which was associated with generally lower antibody concentrations in cord blood, was associated with infection incidence in the first year of life.

**Figure 5:**
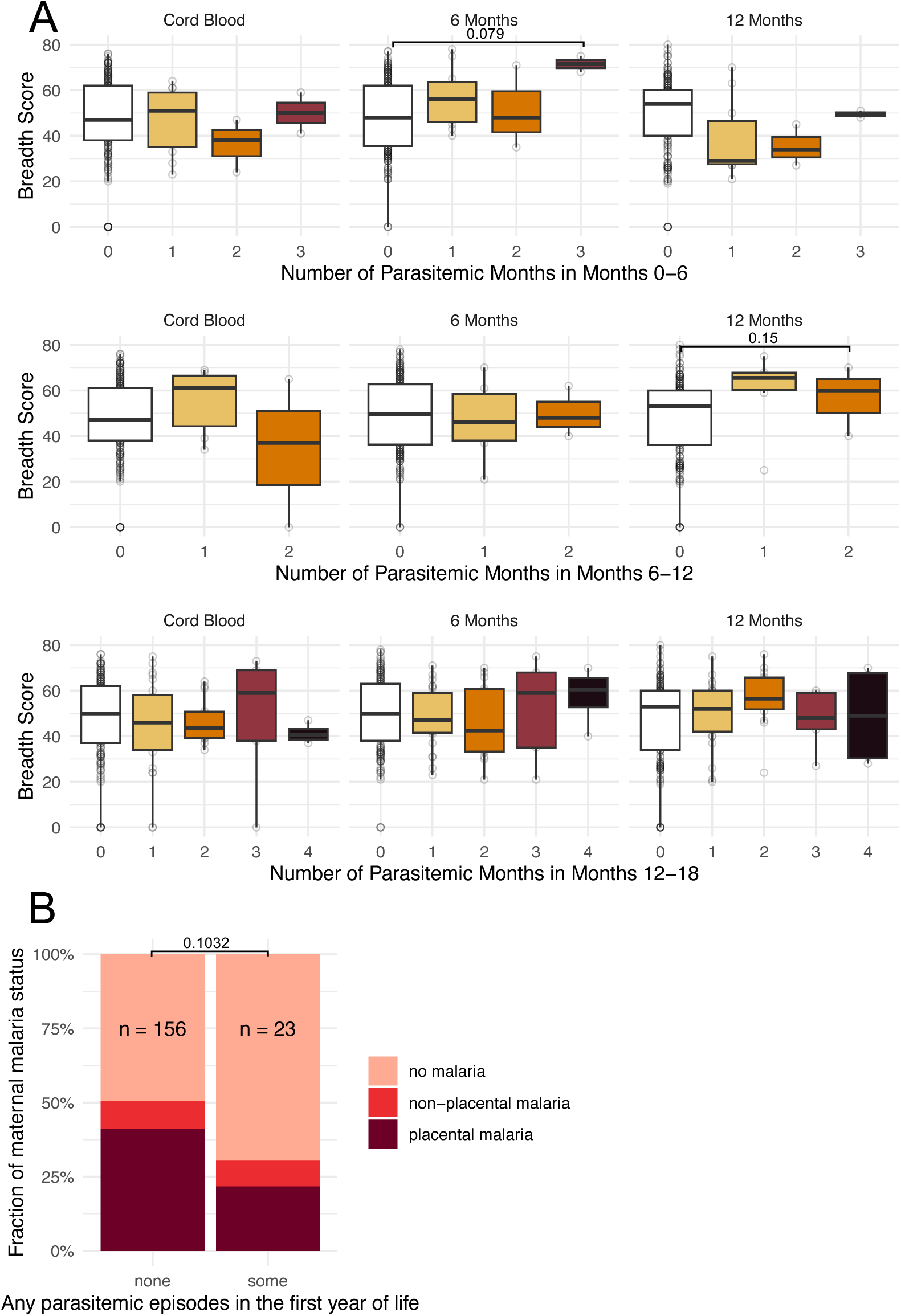
Antibody breadth is associated with exposure, but not protection in early life. (A) Antibody breadth scores at each timepoint were used to model infection incidence in six month increments, as indicated on the x axis. P values shown are BH-adjusted p values of Poisson regression coefficients. (B) Incidence of any parasitemia in the first year of life stratified by maternal malaria status. P value from χ^2^ test.

Individuals born to mothers with placental malaria were not more likely to be infected in the first year of life, despite lower antibody levels in cord blood. In fact, they appeared to experience a lower rate of infection, though this was not statistically significant.

In summary, we found antibodies to several *P. falciparum* antigens were markers of exposure, while none were associated with lower incidence of malaria or prevalent infection. Likewise, the breadth of the repertoire was associated with exposure, not protection. Placental malaria was not associated with early-life prevalence of infection.

### Early life infection is associated with lower antibody levels at 12 months

Next, we considered whether early life malaria, under “cover” of maternal antibodies, may have consequences for the development of acquired immunity during infancy. We found that both the number of parasitemic months and incidence of malaria in the first 6 months of life was associated with significantly lower antibody levels to many *P. falciparum* antigens at month 12 (**Fig. 6**). A possible explanation for this phenotype could be that early-life malaria or asymptomatic parasite carriage negatively impact B and/or T cell development. This prompted us to investigate whether lymphocyte phenotypes at one year differ between children with different early-life infection histories.

**Figure 6:**
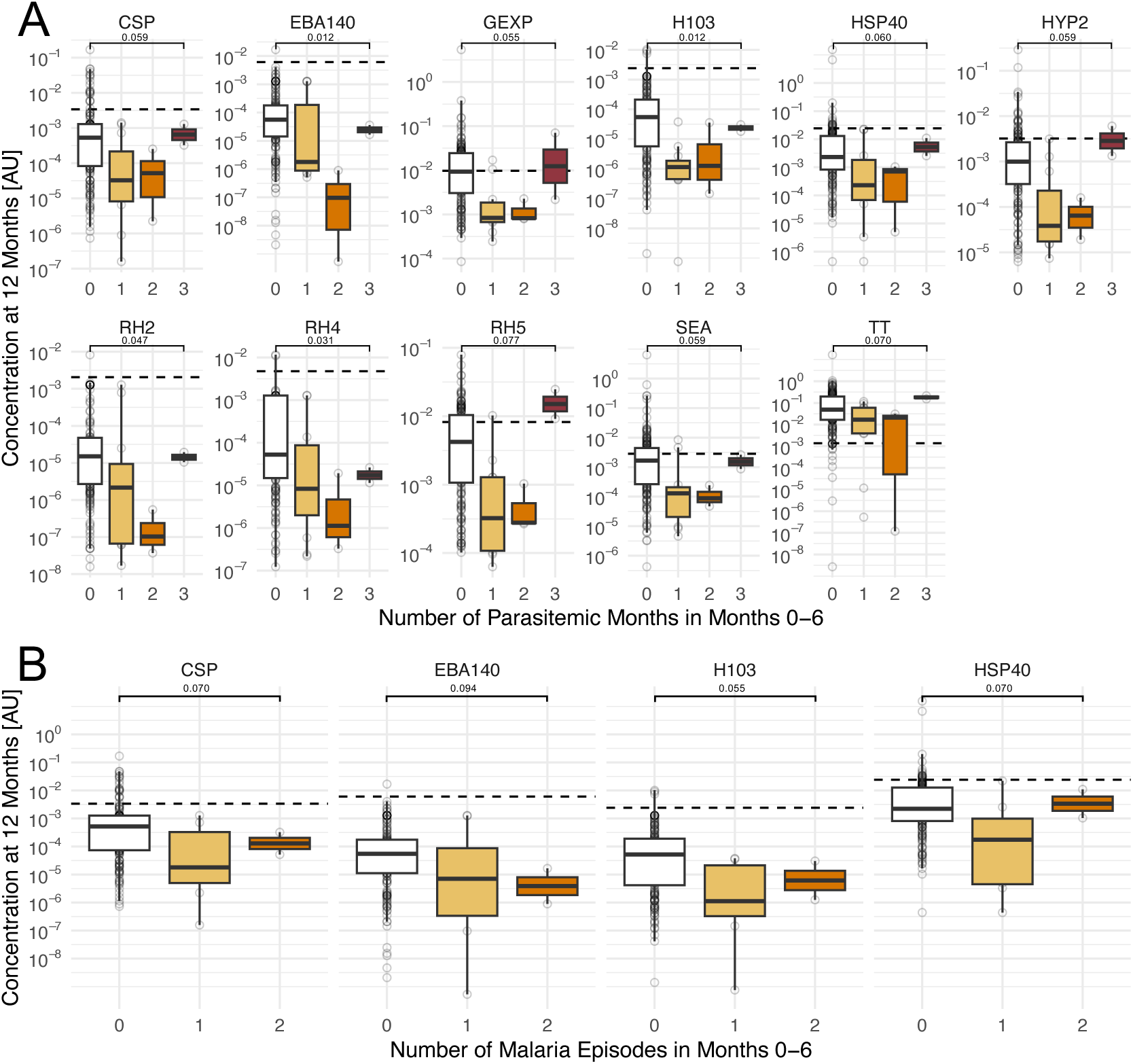
Early life infection is associated with lower antibody levels to *Plasmodium* antigens at one year of age. Both symptomatic malaria **(A)** and number of parasitemic months **(B)** in the first 6 months of life were associated with lower levels of antibodies at one year of age.

### No evidence for major phenotypic alterations in lymphocytes of children exposed to early-life *P. falciparum* infection

To further investigate the mechanisms underlying reduced humoral responses in children exposed to early-life malaria, we decided to phenotypically profile lymphocyte populations important for antibody production, including circulating B cell as well as T helper cell subsets.

We hypothesized that differences in abundance of atypical B cells^7^ and/or circulating follicular T cells (cTfh)^19^ may be associated with altered antibody production.

To determine whether specific T or B cell subsets are associated with malaria infections, we stained fresh PBMCs from 99 and 93 children respectively, to resolve major subsets (see gating strategies in **Fig. S6** and **S7**). We then modelled the incidence of malaria and prevalent infections in the first year of life, based on the frequencies of different lymphocyte populations. (**Fig. 7**, also). We did not find significant associations between early life exposure and abundance of T memory or cTfh phenotypes (**Fig. 7A**). Similarly, neither atypical B cells, nor any other B cell subset was significantly associated with early-life infection incidence (**Fig. 7B**).

**Figure 7:**
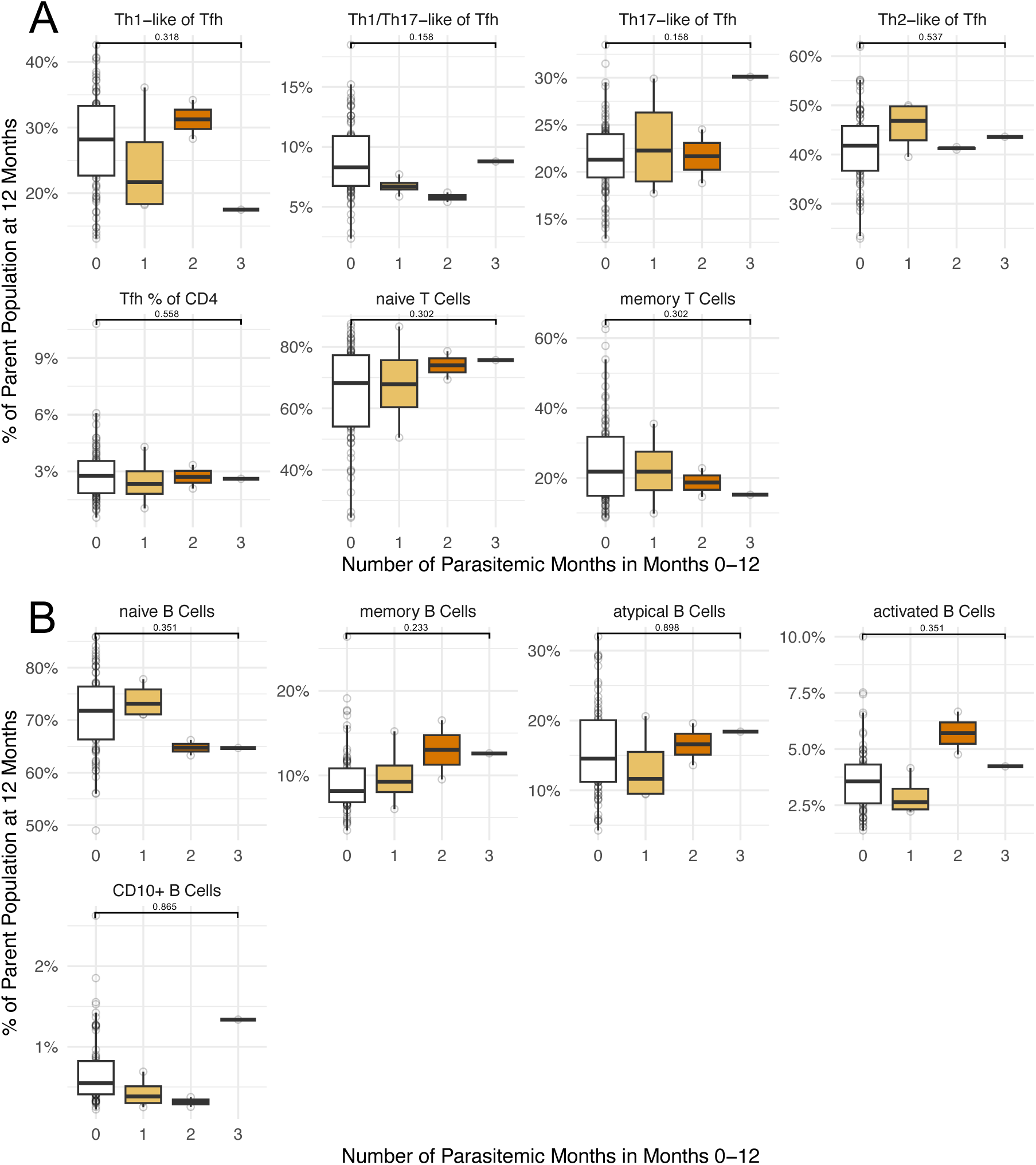
The early life infections are not associated with subset abundance of major lymphocyte populations. Top row: T cell subsets. Bottom row: B cell subsets. P values shown are BH-adjusted p values of Poisson regression coefficients. Gating strategies are shown in figures **S6** and **S7**.

In summary, we found that early-life infections with *P. falciparum* were associated with lower antibody titers at 12 months of age, suggesting some defect in humoral associated with exposure. We tested whether infections during infancy were associated with changes in major B and T cell subsets but did not find significant relationships.

## Discussion

In this study we examined the associations of malaria outcomes during pregnancy and infancy with humoral responses and adaptive immune cell phenotypes. Antibodies were highest in cord blood and decreased quickly during the first 6 months of life whether children experienced infections in that time or not. We found that shorter gestation was associated with reduced levels of antibody transfer, consistent with previous obversations^17,20^. Antibody transfer across the placenta occurs throughout pregnancy, but is increased late in gestation, explaining this finding. We also found consistently lower antibody levels in the cord blood of children born to mothers with placental malaria. While some studies found no association^14,21^, several others identified a significant link between placental malaria and reduced transfer or expression of maternal IgG from the placenta to cord blood^13,23,24^. Placental malaria causes increased rates of birth complications and low birth weight, because fibrosis and inflammation associated with placental malaria limit nutrient transfer^25^. The same tissue pathologies may restrict the active process of IgG transfer to the fetus.

We did not find significant associations between cord blood antibody concentration, that is, maternally transferred antibodies, and reduced incidence of malaria or prevalent infections in early life. Maternally derived antibodies are thought to offer protection against infection and/or disease in infants; however, associations with malaria incidence have been inconsistent^26^. In our study, neither individual antibody concentrations nor repertoire breadth in cord blood were associated with protection. The low burden of infection and lack of complicated malaria in our study means that we may have missed potentially protective properties of these antibodies.

We observed that antibodies to several *P. falciparum protein*s were elevated at 6 and 12 months in children who experienced more infections in the preceding 6-month period. Despite this immune response, these antibodies did not correlate with protection in our study, in contrast to prior work where such associations were found^5,27–29^. Some reasons why antibody concentrations have been inconsistently associated with protection in the literature are discussed below.

An intrinsic problem in the study of humoral immunity to malaria is that antibody concentrations increase with exposure, meaning they are indicative of the risk of exposure, a highly diverse individual and household-level trait that also varies through time^30–32^. Testing whether an antibody might provide protection, that is, decrease the incidence of infection and/or disease, thus ideally involves estimating the individual or household level risk, for example by active case finding over a significant period time. This is often not practical, limiting the utility of studies that lack this information. In studies with large sample sizes and/or homogenous transmission antibodies can be associated with protection, but may not be, possibly explaining some of the heterogeneity in the literature regarding the protective capacity of antibodies. For example, association between SEA antibodies and lower incidence of severe malaria has been described before^33^, however, these antibodies have been associated with increased rates of infection as well^14^, including our study. Lastly, whether antibodies associated with protection are causally responsible, rather than simply correlates of previous exposure that led to protection via other adaptive immune responses is unclear.

Most antibodies provide limited anti-parasite immunity, as evidenced by their often modest growth inhibition *in vitro*, even in highly exposed individuals^34,35^. However, antibodies could mediate protection via mechanisms dependent on immune cells and thus may need to be studied *in vivo*. A seminal set of controlled human infection trials in Kenya ^36^, where potent anti-parasite immunity was observed in some highly-exposed adult volunteers, provided a unique opportunity to study this in humans. Here, above median IgG recognizing schizont extract explained a mere 17% of the variance in time-to-treatment. In an equivalent analysis of a field study with children aged 1-8 years, that number shrunk to 0.8% ^37^. So the amount of antigen-specific antibody may be a poor predictor of protection, as indeed was the case in our study.

Binding capacity is only one facet by of antibody function. Immunoglobulin subtype or post-translational Fc-domain modifications may help mediate protection, by triggering differential cellular responses downstream of Fc-receptor binding^38^. Indeed, Nkumama and colleagues recently reported significant associations between several Fc-mediated antibody functions and clinical immunity during controlled infection^6^. This type of functional characterization, rather than simply measuring antibody abundance, is the way forward to understanding the role of antibodies in antimalarial immunity and is the subject of ongoing studies by us and other groups.

Lastly, we found that *P. falciparum* infection and malaria in the first 6 months of life both were linked to reduced antibody levels at 12 months of age. This is consistent with data showing that high maternal antibodies can lead to decreased post-vaccination titers to many childhood vaccines^39^. The mechanisms underpinning these observations include epitope masking, more rapid clearance of antigen and inhibition of the antigen-specific B cell activation via inhibitory Fc-receptor signaling^40^. This suggests that children living in high-transmission areas, where first-in-life infections happen at a younger age, may develop antibody repertoires more slowly, possibly extending their period of susceptibility.

This defect in antibody production prompted us to query the phenotypic composition of peripheral cell types relevant for antibody production. Unexpectedly, early life exposure did not significantly affect atypical B cell abundance. Studies on B cells in individuals from malaria regions of endemicity demonstrate that *P. falciparum* exposure leads to increased abundance of atypical Memory B cells^41,42^. This was not the case in our study, perhaps because incidence of malaria overall was low, and atypical B cell frequency was more impacted by non-malarial exposures. Instead, we found that children with malaria in the first year of life had somewhat fewer immature (CD10+) B cells and tended to have higher levels of memory B cells compared to uninfected children, but these differences were small and not statistically significant. Likewise, there were no significant associations between infection incidence and T helper subset frequencies.

A key limitation of our study was the low malaria incidence in this cohort, which can be attributed to the implementation of Indoor Residual Spraying (IRS) and effective chemoprevention. It is possible, even likely, that higher infection rates could have revealed more protective associations with the antibodies and cellular phenotypes measured here. While monthly case finding using LAMP means that we likely detected almost all infections, it is possible that very brief infections, or those with parasitemia below the limit of detection went unnoticed. Additionally, the T and B cell assay only queried phenotype and the functional capacity of cells was not assessed. Lastly, our study did not account for potential confounding factors from other infections.

In summary, we found that placental malaria was associated with reduced antibody transfer, as was low gestational age. *P. falciparum*-specific cord blood antibodies were not associated with protection. Similarly, peripheral blood *P. falciparum*-specific IgG obtained at 6 and 12 months of age were associated only with prior malaria exposure, not protection. Surprisingly, we found that early-life malaria was associated with decreased antibody responses at one year of age, which was not explained by overt differences in peripheral B and T cell subset composition.

## Materials and Methods

### Study design

The objective of this study was to evaluate antibody and cellular immune responses in cord and peripheral blood of infants born to mothers living in a high malaria transmission setting. Of the 300 pregnant women enrolled in the parent clinical trial, 291 were followed until delivery. Of the infants born to these mothers, a subset (186 for antibody responses and a partially overlapping n = 102 for cellular responses, see supplementary figure 3) was included in this study. Serum for antibody responses was collected at birth, 6 months and 12 months and stored at −80°C. Peripheral blood mononuclear cells were isolated from whole blood collected at one year using Ficoll gradient centrifugation using standard protocols and used fresh for assessing cellular phenotypes by flow cytometry.

### Ethical approval

Informed consent was obtained from the mothers of all study participants. The study protocol was approved by the Uganda National Council of Science and Technology and the institutional review boards of the University of California, San Francisco, Makerere University, and the Center for Disease Control and Prevention.

### Study site and participants

This study was nested within a parent trial, the PROMOTE II Birth Cohort I (BC1) study carried out in Tororo, Uganda. The parent study was a double-blind randomized placebo-controlled phase III trial, which involved 300 HIV uninfected pregnant women and the children born to them. Participants at 12-20 weeks’ gestation were randomized in equal proportions to one of the three intermittent preventive treatment in pregnancy (IPTp) arms: a) three doses of sulfadoxine pyrimethamine (SP, standard of care), b) 3 doses of dihydroartemisinin piperaquine (DP), or c) monthly DP. All three interventions arms had either SP or DP placebo to ensure that adequate blinding was achieved in the study. At birth, cord blood and placental tissue were assessed for the presence of malaria parasite DNA by loop-mediated isothermal amplification (LAMP) and for histopathologic evidence of placental malaria (PM), as previously described^43,44^. All children born to mothers enrolled in the study were followed from birth until they reached 36 months of age. Children born to mothers randomized to receive 3 doses of SP during pregnancy received DP every 3 months between 2-24 months of age. Children born to mothers randomized to receive 3 doses of DP or monthly DP during pregnancy received either DP every 3 months or monthly DP between 2-24 months of age. All but one child in this study received DP every 3 months. Children were then followed up for an additional year between 24-36 months of age following the cessation of interventions^44^.

### Screening for maternal parasitemia during pregnancy

Maternal parasitemia was assessed every 4 weeks throughout pregnancy, beginning at enrollment. The presence of *Plasmodium spp.* parasites in maternal peripheral blood was evaluated using LAMP kits (Eiken Chemical), as previously described. LAMP assays were analyzed at the end of the trial and were not used to inform treatment (asymptomatic parasitemia is not treated per current Ugandan standard of care). Placental tissue was processed for histopathologic evidence of placental malaria, as previously described^45^, and placental blood and cord blood were tested for the presence of *Plasmodium* by both LAMP and microscopy.

### Childhood malaria outcomes

After birth, infants were given DP every 4 or 12 weeks until 2 years of age and then followed to 3 years of age. Routine visits were conducted every 4 weeks and included the assessment of peripheral parasitemia by LAMP. Children were encouraged to be brought to the clinic any time they were ill. Those who presented with a documented fever (tympanic temperature > 38.0°C) or history of fever in the previous 24 hours had blood collected for a thick blood smear, and if parasites were present, the patient was diagnosed with clinical malaria. Episodes of uncomplicated malaria in children <4 months of age or weighing <5 kg, as well as episodes of complicated malaria and treatment failure within 14 days, were treated with a 7-day course of quinine.

### Luminex Magpix assay

We utilized a Luminex multiplex bead array to assess total IgG antibody responses to 19 *P. falciparum* antigens and tetanus toxoid, allowing for high throughput, low-cost assessment of several serological markers together (see Table 1). Plasma samples obtained at birth (cord blood), 6 months and 12 months were evaluated. Beads were conjugated with Pf antigens, as previously published^46^. Quantification was performed according to standard protocols. Briefly, duplicate samples of 50 µL thawed plasma were diluted 1:200, co-incubated with microsphere mixtures on a 96-well plate for 1h, washed, stained with an anti-human Fc secondary antibody, then washed and read with a MAGPIX system. Antibody levels were expressed in arbitrary units (AUs), calculated by dividing the median fluorescence intensity (MFI) of the sample by the MFI plus 3 standard deviations (SD) of samples from North American adults never exposed to malaria. Positive control samples from individuals with known antibodies to these antigens were included on each plate. Standard curves were generated through serial dilutions of the positive control pool. To obtain relative concentrations of antibodies and to account for plate-to-plate variation, raw MFI were regressed onto the standard curve for each antigen present on every plate. Seropositivity thresholds are shown as a dashed line in all plots. Our primary analysis evaluated responses to each antigen separately.

### Flow cytometry

6 mL of whole blood were obtained from each infant in acid citrate dextrose tubes. Peripheral blood mononuclear cells (PBMC) were isolated by Ficoll density gradient centrifugation using standard protocols. Fresh PBMC were used in all assays. PBMC were stained with fluorescently conjugated monoclonal antibodies specific for immune cell surface markers and/or chemokine receptor markers that allow for identification of CD4^+^ T cell and B cell memory and lineage subsets as shown here:

**Table 3:** Flow cytometry reagents.

| Antibody | Color | Clone | Source | Cat# |
| --- | --- | --- | --- | --- |
| CXCR3 | BV421 | G025H7 | Biolegend | 353716 |
| CD3 | APC-Cy7 | HIT3a | Biolegend | 300318 |
| PD-1 | FITC | EH12.2H7 | Biolegend | 329936 |
| CD45RA | PerCP-Cy5.5 | HI100 | Biolegend | 304122 |
| CCR6 | PE | G034E3 | Biolegend | 353410 |
| CCR4 | PE-Cy7 | L291H4 | Biolegend | 359410 |
| CXCR5 | APC | J252D4 | Biolegend | 356908 |
| CD4 | BV510 | RM4-5 | Biolegend | 100553 |
| CD27 | BV421 | M-T271 | Biolegend | 356418 |
| CD14 | BV510 | M5E2 | Biolegend | 301842 |
| CD3 | BV510 | UCHT1 | Biolegend | 300448 |
| CD21 | PE | HB5 | eBioscience | 12-0219-42 |
| CD20 | APC-Cy7 | 2H7 | Biolegend | 302314 |
| CD10 | FITC | HI10a | Biolegend | 312208 |

Cell acquisition was done with a FACS CANTO flow cytometer running FACSDiva software. Color compensations were performed using a mix of all patients’ PBMC single-stained for each of the fluorochromes used. Data was then analyzed using FlowJo (African Program). The gating strategy for identification of T and B cell subsets is summarized in supplementary figures 1 and 2, respectively.

### Statistical Analysis

All statistical analysis was carried out in R (version 4.4). Simple unpaired group-comparisons were performed using Wilcoxon rank sum tests or Chi squared tests, as indicated. Associations of immunological assay results with incidence measures were analyzed using linear regression. Incidence measures, the number of infections or malaria episodes in a given time window, were modelled using Poisson regression implemented in *stats* (version 4.4). Separate regression models were fit for each association. Multiple testing correction using Benjamini-Hochberg (BH) was performed on one-sample t tests of regression coefficients, with p values batched by timepoint. Due to high correlation between features of interest (e.g. all malaria-specific antibodies are expected to be potentially boosted by infection), we report results from our exploratory analyses as significant if their BH-adjusted p value < 0.1. We stress that these are exploratory analyses and the low level of malaria transmission in this study may limit the generalizability of our findings.

To calculate an antibody breadth score, each individual was sorted individuals into antigen-specific quartiles of antibody concentration. The quartile positions (1 to 4) were then summed to calculate a breadth score for each individual and each timepoint.

## Supplementary Figures

**Supplementary figure S1:**
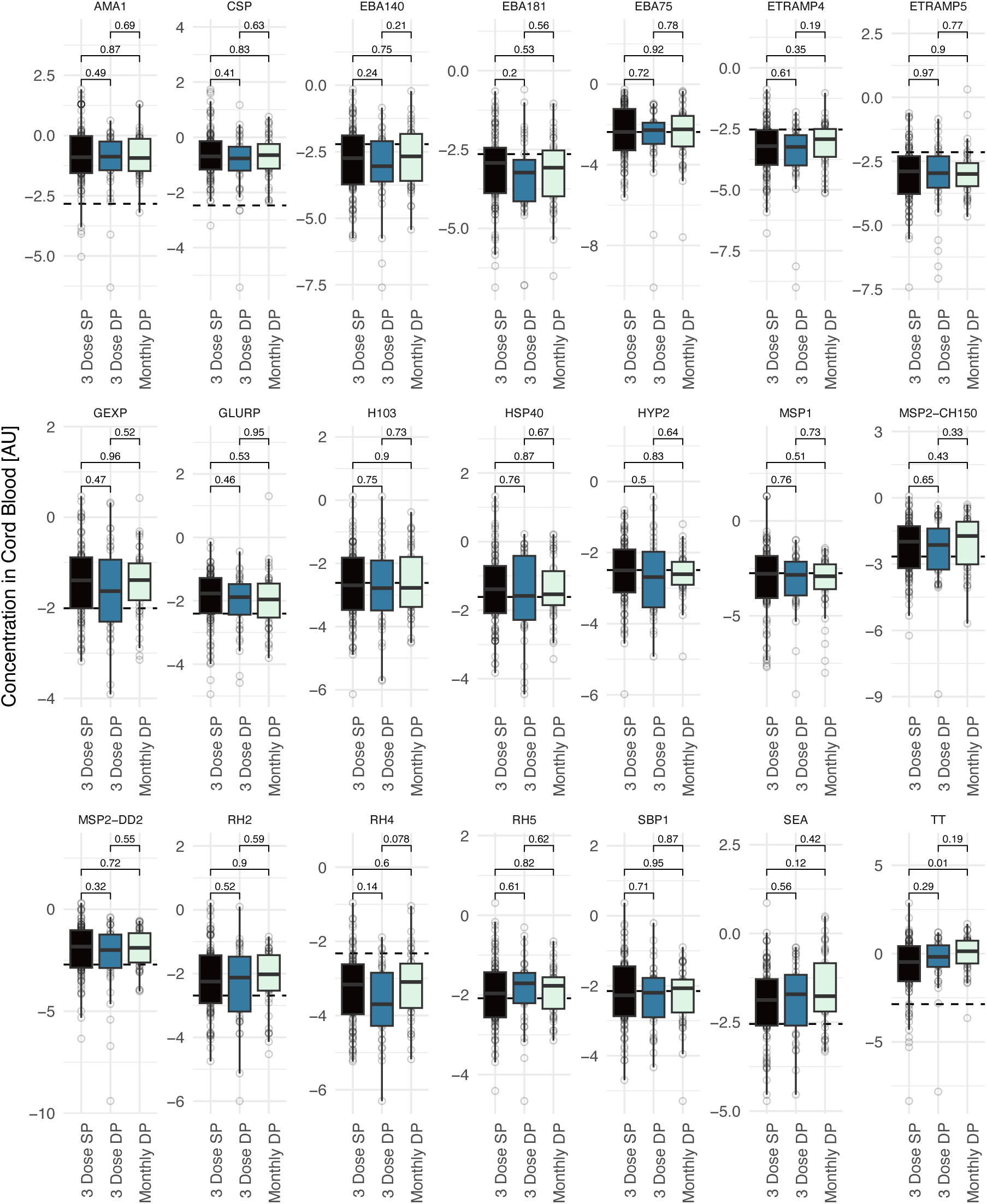
Cord blood antibody concentrations stratified by maternal chemoprevention regimen

**Supplementary figure S2:**
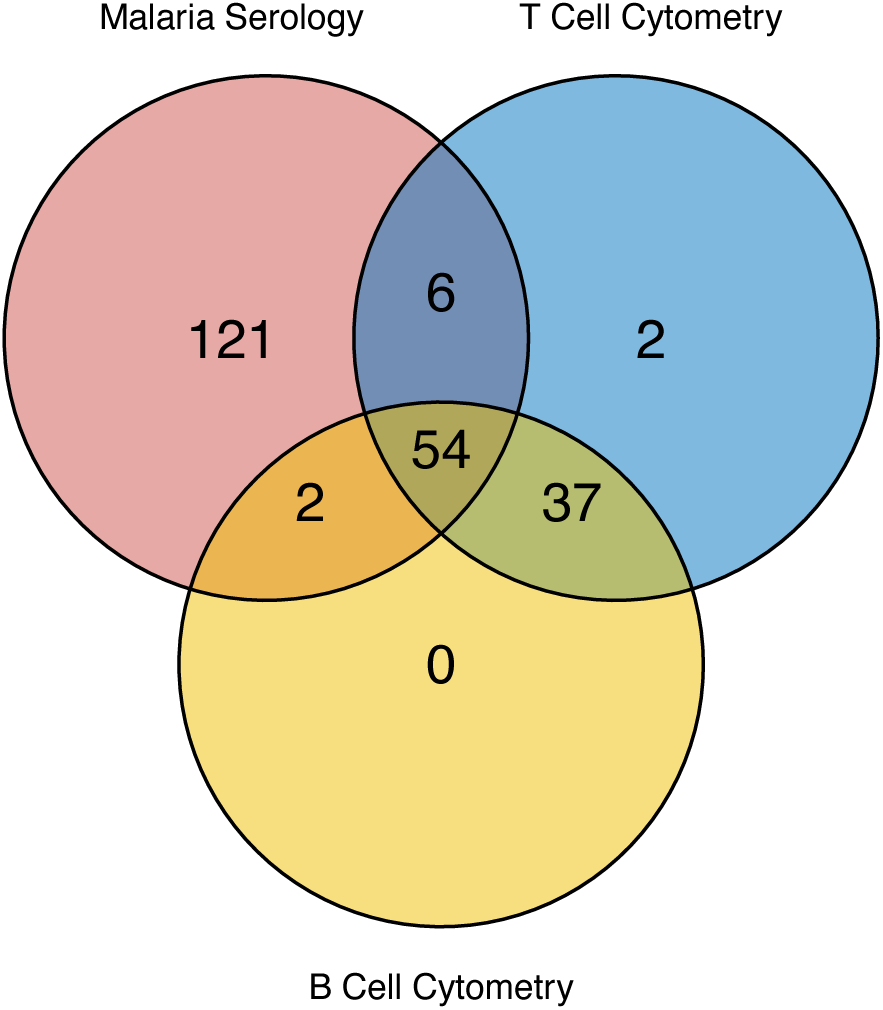
Venn diagram showing how sampling cohorts for serology, T cell and B cell cytometry overlap

**Supplementary figure S3:**
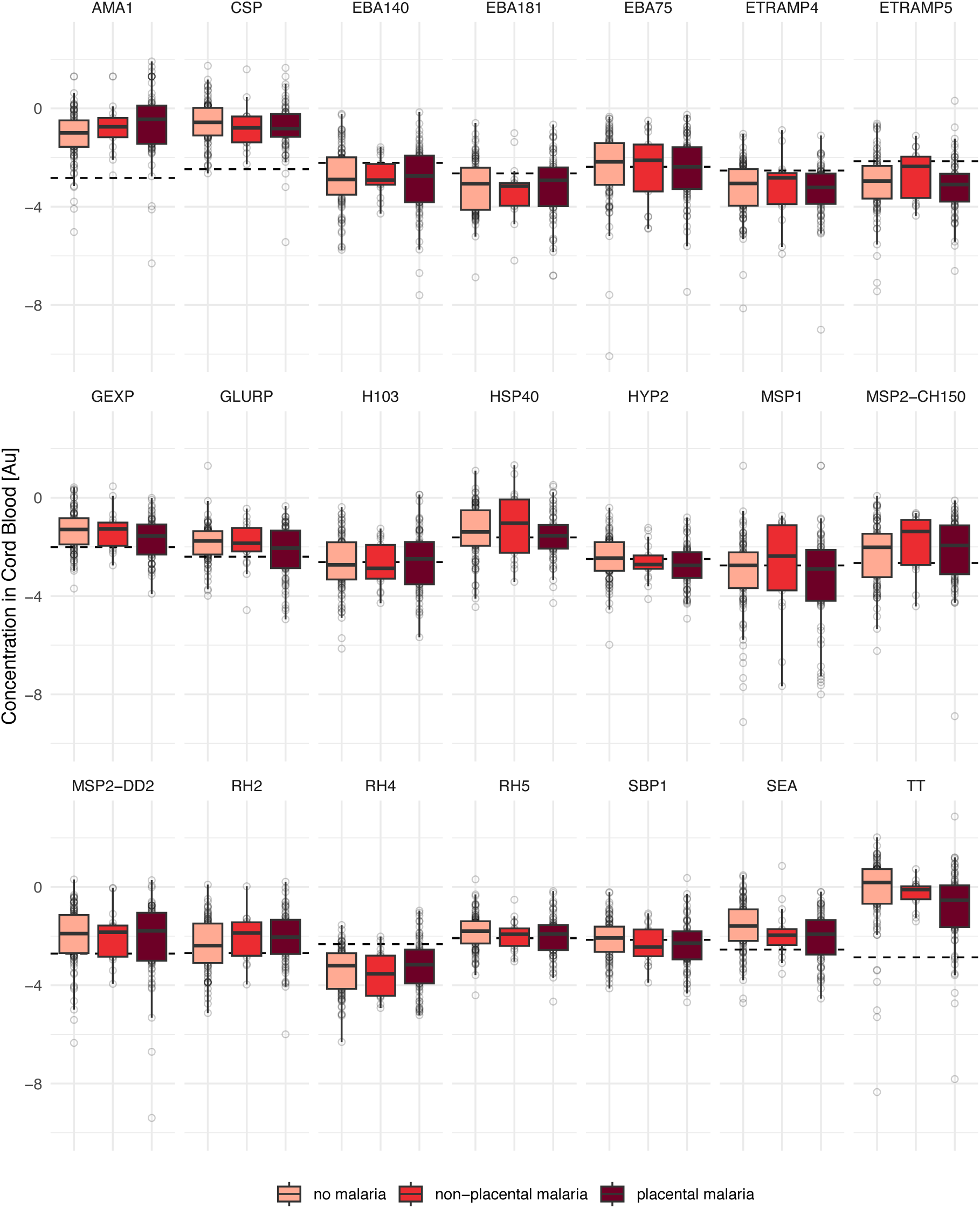
Cord blood antibody concentrations stratified by maternal malaria status during pregnancy

**Supplementary figure S4:**
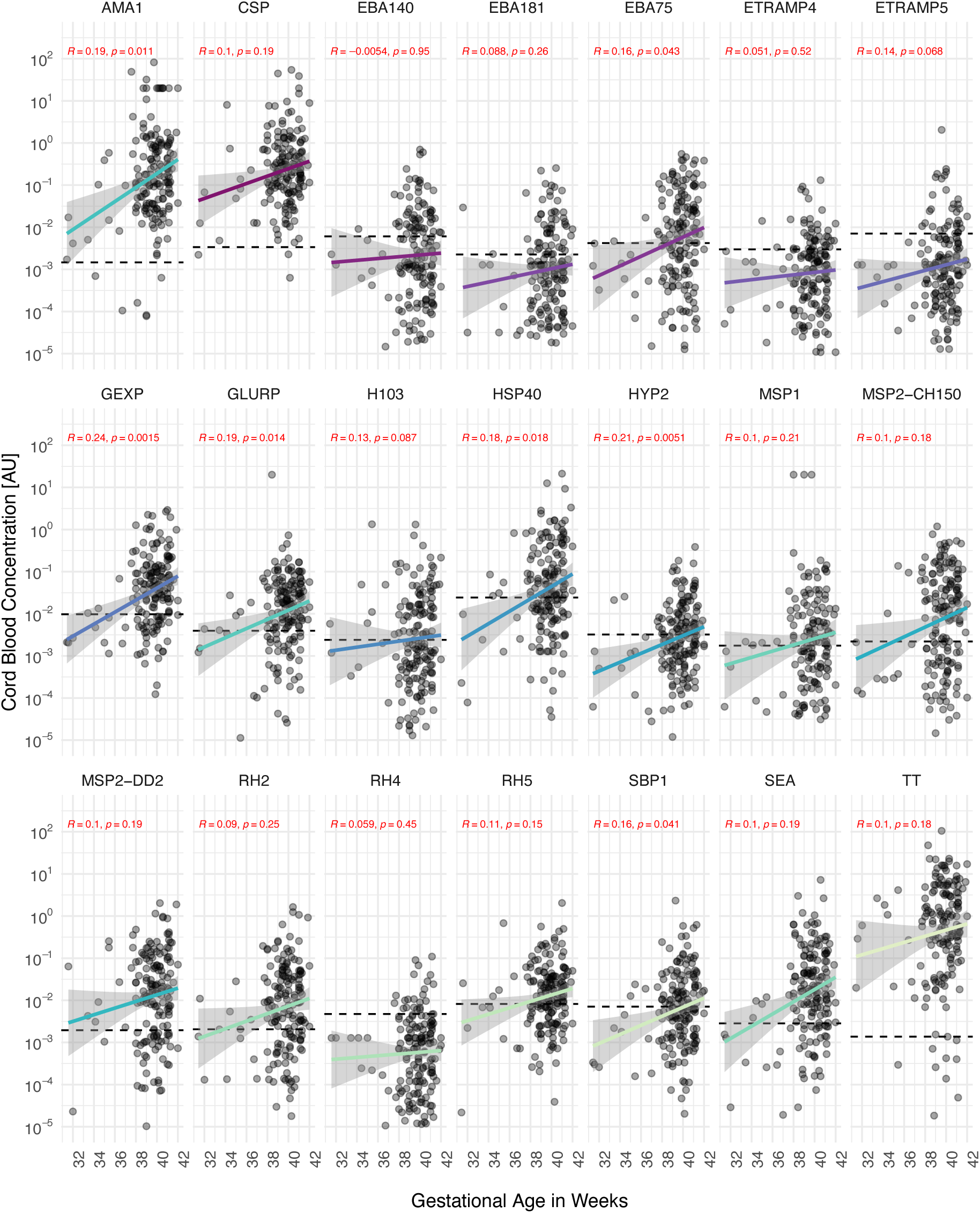
Cord blood antibody concentrations correlated with gestational age of infant

**Supplementary Figure S5:**
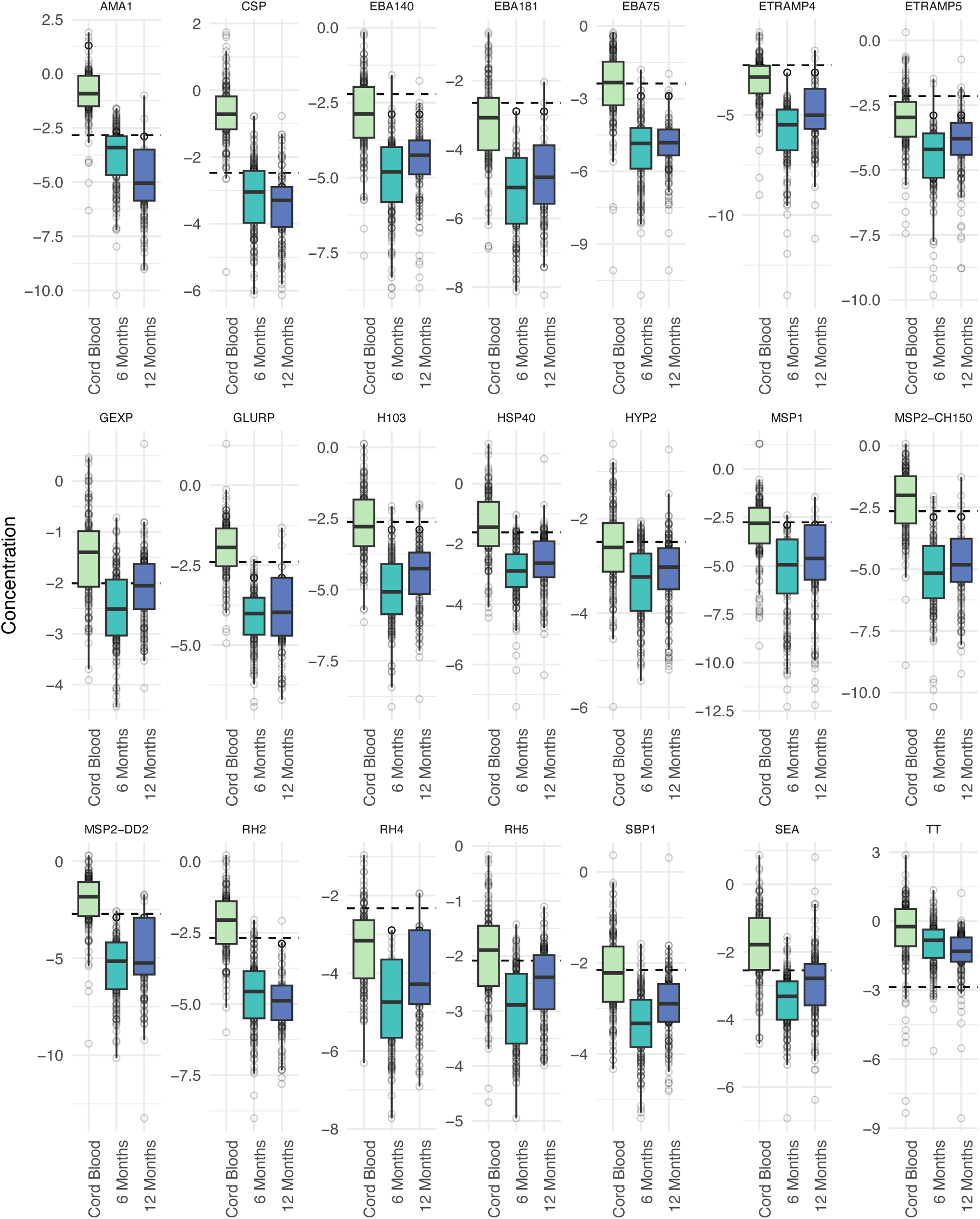
*P. falciparum* antibody kinetics in the first year of life in children who remain uninfected.

**Supplementary Figure S6:**
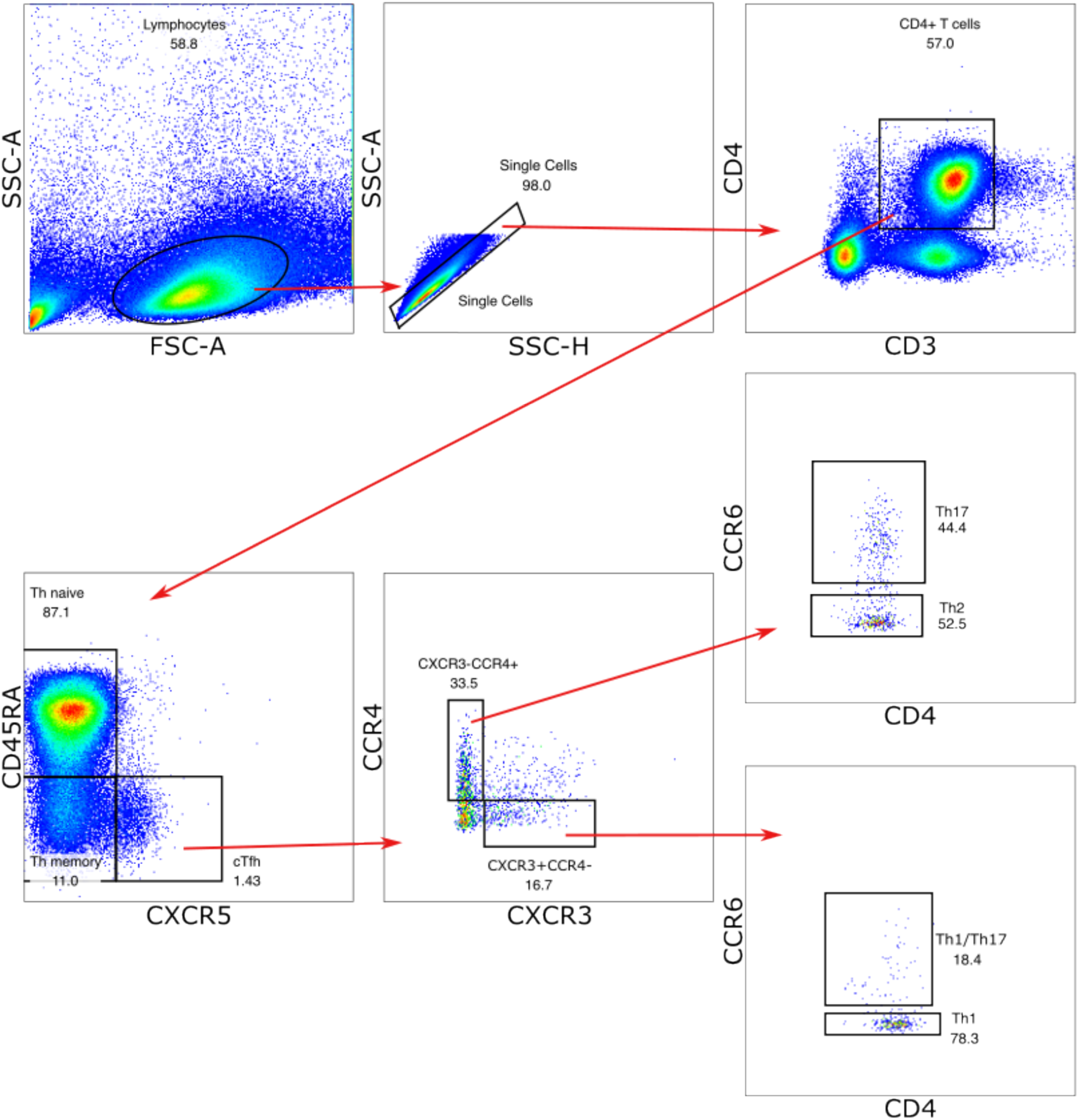
Identification of T cell subsets.

**Supplementary Figure S7:**
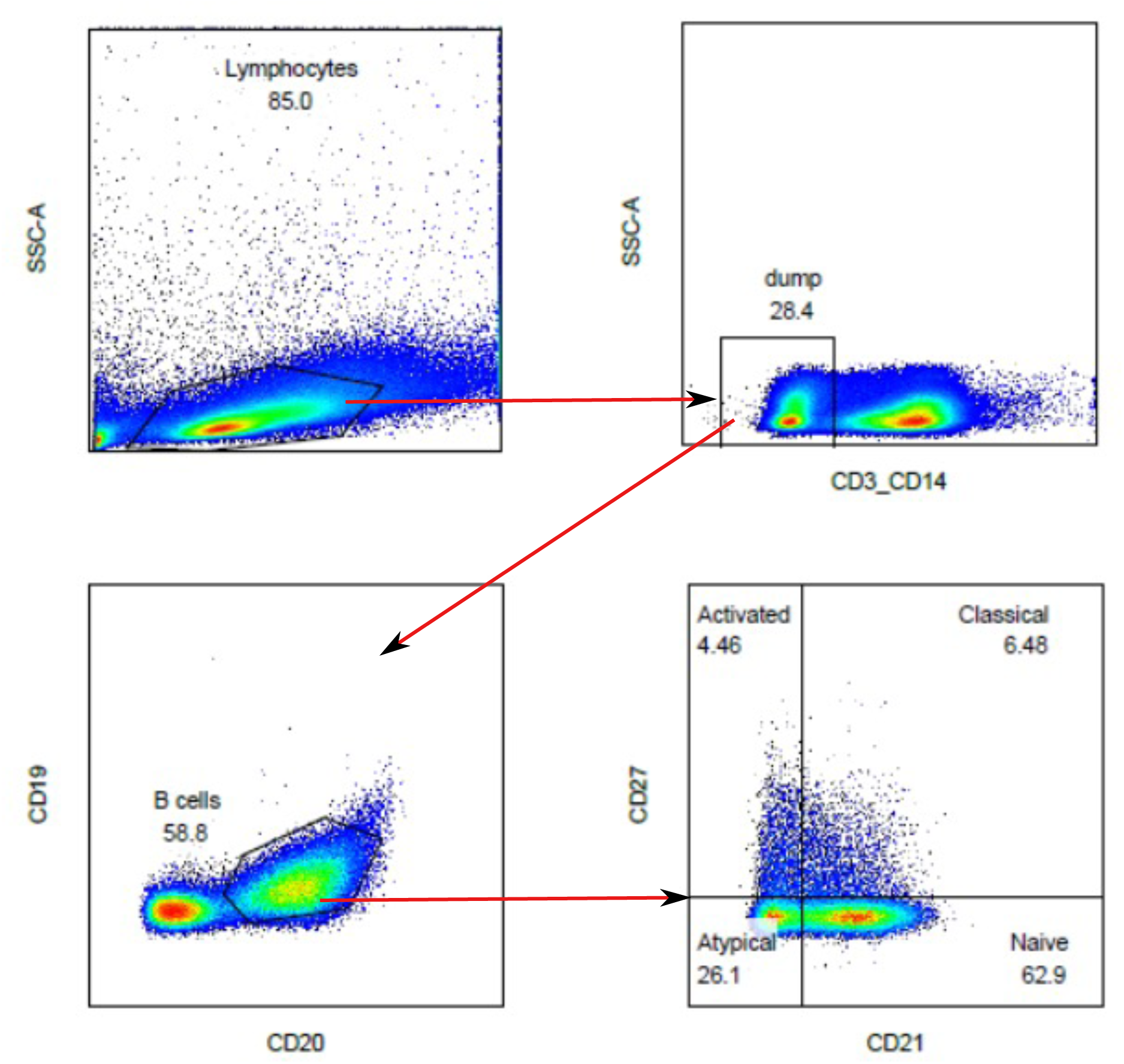
Identification of B cell subsets.

